# MolSnapper: Conditioning Diffusion for Structure Based Drug Design^†^

**DOI:** 10.1101/2024.03.28.586278

**Authors:** Yael Ziv, Brian Marsden, Charlotte M. Deane

## Abstract

Generative models have emerged as potentially powerful methods for molecular design, yet challenges persist in generating molecules that effectively bind to the intended target. The ability to control the design process and incorporate prior knowledge would be highly beneficial for better tailoring molecules to fit specific binding sites. In this paper, we introduce MolSnapper, a novel tool that is able to condition diffusion models for structure-based drug design by seamlessly integrating expert knowledge in the form of 3D pharmacophores. We demonstrate through comprehensive testing on both the CrossDocked and Binding MOAD datasets, that our method generates molecules better tailored to fit a given binding site, achieving high structural and chemical similarity to the original molecules. It also, when compared to alternative methods, yields approximately twice as many valid molecules.

## 1 Introduction

Drug design is a complex process involving iterative optimization steps to achieve desired biological responses. The vast search space and discontinuous nature of the optimization landscape pose significant challenges, which has led to a reliance on human experts for molecular design. Traditional methods, typically based on trial-and-error approaches, result in high costs and limited productivity^1,2^.

Computational methods have long been central to drug discovery, aiming to reduce costs, expedite processes, and minimise failures^3^. The emergence of artificial intelligence, especially deep learning, has shown enormous potential to revolutionise the field, it initially found application in property prediction, yielding promising early results^4,5^. The success of deep learning in this context stems from its flexibility to directly learn from raw data^6,7^. One area of application of deep learning models is compound design. In this area models generate multiple molecules, aiming to suggest those with desirable properties^8–10^. One of the most important properties that has been the subject of this type of work is binding of the molecule to its target^11^.

In the field of structure-based drug design (SBDD), the objective is to generate ligands with high affinity and specificity for a specific protein in a specific 3d conformation. However, designing ligands precisely tailored to bind to a target protein remains a persistent challenge. Several machine learning models have been employed in SBDD, for example, autoregressive models^12–14^. These models generate 3D molecules within the target binding site by iteratively adding atoms and bonds. However, a limitation of autoregressive models is the accumulation of errors during the generation process. Additionally, sequential generation methods may not fully capture the complexities of real-world scenarios, as they impose an artificial ordering scheme, potentially losing the global context of the generated ligands.

To address the limitations of autoregressive models, recent studies^9,15–18^ have turned to diffusion models^19^. These models iteratively denoise data points sampled from a prior distribution to generate samples. Unlike autoregressive models, diffusion model-based methods can simultaneously model local and global interactions between atoms, leading to improved performance^16,20^.

Despite these advances, computational SBDD faces challenges in terms of synthesizability and chemical feasibility of the generated molecules. The limited volume of experimentally determined structures of protein-ligand complexes often leads models to learn dataset biases rather than grasping the true biophysical principles underlying ligand-protein interactions^21^. Diffusion methods have also been developed and trained on the far larger dataset of just molecules which allows better coverage of druglike space potentially leading to more viable and synthesizable drug candidates^15^.

The practicality of computational SBDD methods is also hindered by an inability to explicitly include expert knowledge, a limitation that becomes particularly evident in the later-stages of drug discovery where substantial prior knowledge could and should guide compound design. For deep learning methods to gain widespread adoption, more control over the generative process is essential^22^.

Examples of methods that have used expert knowledge to guide compound design include DEVELOP^22^, PGMG^23^ and STRIFE^24^, which employ autoencoder architectures. These methods, which generate either 1D smiles strings or 2D graphs of molecules, integrate target pharmacophores into the generative process, leading to enhanced molecule generation with superior control over the design phase. The authors show how the integration of pharmacophoric constraints significantly improves the quality of the molecules generated, via multiple large-scale evaluations.

Conditioning, which involves providing additional information or constraints to a model during generation, can be also used to achieve control. Pre-trained diffusion models represent one approach to delivering this control. Methods exist that use conditioning for molecule generation in a protein pocket, for example, SILVR^25^ (Selective Iterative Latent Variable Refinement). This method conditions existing diffusion-based models for the generation of new molecules that fit into a binding site without knowledge of the protein. SILVR introduces a refinement step within the denoising process, employing a linear combination of the generated ligand and the noised reference. However, it requires an equal number of reference and generated atoms, focusing primarily on fragment merging and linker generation. Due to the lack of any knowledge about the protein, the presence of several dummy atoms can lead to clashes of the generated molecules with the protein^26,27^.

Another conditioning method inpainting is used by DiffSBDD^16^. In inpainting, an unconditional diffusion model can generate approximate conditional samples when the context is injected into the sampling process by modifying the probabilistic transition steps. DiffSBDD, directly incorporates protein information into the model (rather than in the conditioning process), yet it retains the restriction of using only protein-ligand complex data, in its training.

Recent studies PoseBusters^26^ and PoseCheck^27^ have highlighted that deep learning methods in SBDD, including diffusion models, frequently produce molecular structures that are physically implausible. This problem arises partly because the models’ performance metrics are not aligned with physical viability. Specifically, deep learning generative models struggle to create the hydrogen bond interactions between the target protein and ligand seen in the ground truth ligands. PoseCheck found that none of the seven methods tested matched or exceeded the ground truth ligand’s interactions with its protein target and that the most common number of hydrogen bond acceptors and donors of the generated molecules was zero.

Moreover, the evaluation of SBDD-generated poses often includes a redocking step, which can mask issues such as steric clashes and elevated strain energies^27^. Essentially, for a model to be practically useful, its generated molecules should be viable without the need for redocking, which can significantly shift the molecules from their model-generated positions (PoseCheck demonstrated an average RMSD of 0.94 to 1.28 Å between the redocked and original poses).

In this paper, we propose MolSnapper, a novel tool that conditions diffusion models for SBDD by integrating expert knowledge. Our approach is focused on generating molecules that are not only plausible and valid but also capable of forming hydrogen bonds or other interactions similar to those observed in ground truth ligands. While we utilize physically-meaningful 3D structural information, typically provided as 3D pharmacophores^28^, our framework is not limited to this type of constraint. Recognizing the diverse nature of protein targets, MolSnapper allows users the flexibility to employ various constraint types tailored to their specific protein of interest. These constraints can be manually selected or automatically extracted from experimental data, such as fragment screening experiments.

Protein information is leveraged in the form of user guidance, offering a more adaptable framework for drug design and allowing the utilization of models trained on large molecule datasets for molecule generation. This contrasts with SBDD models that are often trained on limited protein-molecule data.

We have benchmarked MolSnapper using the CrossDocked^29^ and Binding MOAD^30^ datasets. In our evaluation, MolSnapper outperforms the SILVR approach, which similarly to our approach, operates without additional training on protein-ligand data. MolSnapper also achieves results comparable to dataset-specific conditioning methods trained on specific protein-ligand data. Our results demonstrate the efficacy of MolSnapper in improving the proportion of generated molecules that closely resemble the reference molecule. Furthermore, our approach successfully reproduces the majority of hydrogen bond interactions observed between the reference ligand and the protein and yields approximately twice as many valid molecules as other methods.

## 2 Method

We create a generative diffusion model to predict molecules for a given protein pocket that takes advantage of the large available training data in drug-like molecule space and integrates physically meaningful 3D structural information to the generative process using 3D pharmacophores^28^.

As our base model we use MolDiff^15^ as it is among the top-performing models for molecule generation^15^. MolDiff was pretrained on the GEOM-Drug dataset^31^. We do not retrain MolDiff; instead we condition it without altering the model weights.

A 3D small molecule is characterized by atom types, atom positions (coordinates in space), and bonds type. A molecule with N atoms can be denoted as *M* = {*A, R, B*}, where *A* = {*a*_*i*_}_*N*_ *∈***A**^*N*^ represents atom types, *R* = {*r*_*i*_}_*N*_ *∈***R**^*N×*3^ represents atom positions, and *B* = {*b*_*i j*_}_*N×N*_ *∈***B**^*N×N*^ represents chemical bonds. Additionally, specific positions related to pharmacophores are denoted as 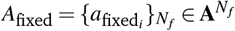, representing the fixed positions and types of the pharmacophores, and 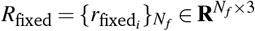, representing the fixed atom positions associated with pharmacophores, and 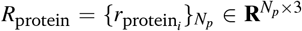, representing the positions of the target protein.

In the context of the diffusion model framework, following the framework introduced in the MolDiff paper^15^, during the reverse process, the Markov chain is reversed to reconstruct the true sample. This involves using *E*(3)-equivariant neural networks to parameterize the transition *p*_*θ*_ (*M*^*t*−1^|*M*^*t*^) from prior distributions, where 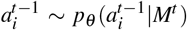 and 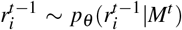. Specifically, the predicted atom positions are modeled as a Gaussian distribution 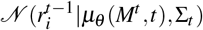, and the atom types are modeled as Categorical distributions 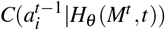 where μ_*θ*_ and *H*_*θ*_ are neural networks. Here, Σ_*t*_ is set as *β*_*t*_ where *β*_*t*_, *t ∈* [0, 1] is the pre-defined noise scaling schedule.

In our approach, the unconstrained diffusion process is altered in that the parameterized distribution is conditioned by the pharmacophores and the protein positions. We modify the reverse process such that 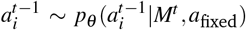 and 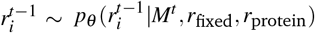, while the bonds prediction remains unchanged.

### 2.1 Pharmacophores

Pharmacophores can be used as a representation of chemical interactions crucial for ligand binding to macromolecular targets^28^. These interactions include hydrogen bonds, charge interactions, and lipophilic contacts. Pharmacophores can be derived through ligand-based or structure-based approaches, providing versatility in their application. In this study, we employed 3D pharmacophores derived from ground truth molecules, using the 3D positions and types (donor or acceptor) of these pharmacophores to condition the reverse generative process.

Given their widespread relevance in drug discovery, our chosen pharmacophores included hydrogen bond donor and acceptor features, determined according to default RDKit definitions. Hydrogen bonds both play an important role in various proteinligand interactions^32^, and, generative models, as highlighted by PoseCheck^27^, often face challenges in reproducing hydrogen bond interactions compared to the ground truth ligand.

While our framework incorporates specific constraints in this study, it is inherently flexible and can be extended to include additional constraints based on user preferences. Users can tailor the features to guide the design process according to their specific requirements.

### 2.2 Constrain Positions

To constrain the reverse generation of atom positions, a mask *X ∈* {0, 1} is introduced, where Mask *X* is applied to limit the modification to the positions of the pharmacophores. The pre-defined noise scaling schedule *β*^*t*^ is incorporated:

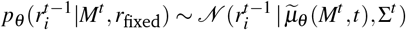

Where:

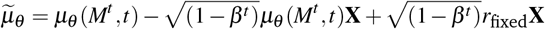

At each stage of the process, we compel certain atom positions to approach the fixed locations. In the early steps, we facilitate the atoms to move slightly toward the fixed positions, while in the final steps, we firmly anchor them in place.

Unlike other methods, we adopt the strategy of utilizing the original positions of the pharmacophores reference points without introducing additional noise. This decision is grounded in the observation that in diffusion models for molecule generation, we operate the reverse process in the same space as the molecule space. Our empirical findings indicate that this approach yields comparable or even superior results compared to incorporating additional noise as can be seen in the results section. Furthermore, rather than opting for random initialization, we initialize the positions randomly but around the pharmacophore positions. In Figure 1, we illustrate sampling of positions constrained by fixed positions.

**Fig. 1.**
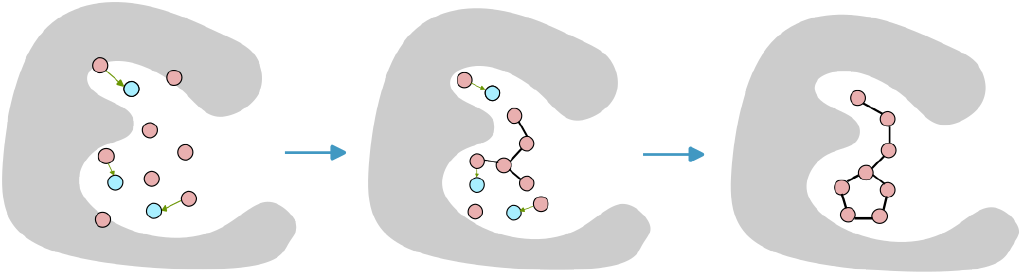
Visualization of the sampling process depicting atom positions constrained by fixed locations. The pink spheres represent the positions of the generated molecule, the blue spheres indicate the positions of the fixed reference points, and the gray mesh represents the protein surface. At each step after the diffusion outputs the position at time *t*, some atoms are gradually moved towards the fixed positions, aiding in the refinement of the molecular structure.

### 2.3 Guidance Preventing Clashes

Since the MolDiff model was not trained with protein data, it cannot process pocket information. Therefore, it is crucial to incorporate guidance to prevent clashes or excessively short distances between the generated ligand and the protein. As proposed by Sverrisson et al.^33^ and Guan et al.^20^, we describe the protein’s surface as

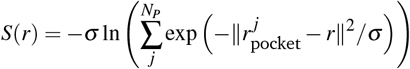

Where 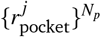 represent the protein atoms positions. following Guan et al.^20^, the clash guidance loss function is defined as:

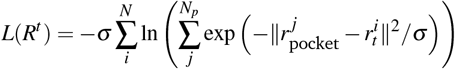

Then the gradient of *L*(*R*^*t*^) with respect to *R*^*t*^, (∇*L*(*R*^*t*^)) provides the direction to enhance molecule generation. With the guidance, the reverse generation of the atom positions

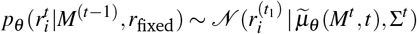

can be adjusted as

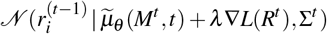

where *λ >* 0 is a constant coefficient controlling the strength of the guidance.

Figure 2 illustrates the clash guidance.

**Fig. 2.**
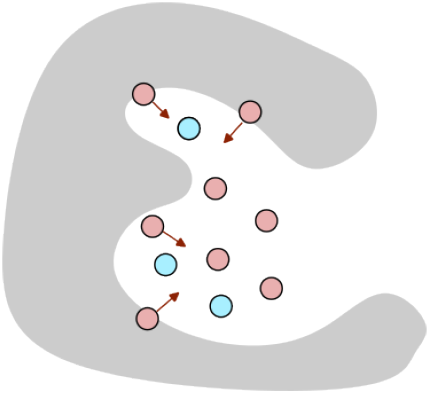
Illustration of clash guidance in the molecule generation process. The fixed positions are shown in blue, the pink positions represent the generated molecule, and the gray mesh represents the protein surface. Clash guidance, inspired by Guan et al.^20^, prevents clashes or excessively short distances between the generated ligand and the protein. The gradient of the clash guidance loss, denoted as ∇*L*(*R*^*t*^), provides the direction for enhancing molecule generation.

### 2.4 Constrain Atom Types

To enhance interpretability and control over the generated molecules, we allow users to constrain atom types based on their specific requirements. Users can specify the desired atom types in the reference, such as based on the interactions they wish to mimic or the reference structure they intend to use. Throughout the iterative sampling process, the model consistently applies these user-defined constraints, ensuring that the generated molecules align with the intended design objectives. This approach provides flexibility for domain experts to modify this choice as needed in the final output.

Our method introduces a hyperparameter representing the Pharmacophore Satisfaction threshold, evaluating whether the ligand fulfilled the 3D pharmacophore constraints (distance lower than 1Å and matching family type). We kept only those ligands meeting this criterion across three attempts.

#### Code availability

The code is available at https://github.com/oxpig/MolSnapper.

## 3 Experiments

### 3.1 Experimental Setup

#### 3.1.1 Datasets

We evaluate the efficacy of our conditioning method on two datasets: the synthetic CrossDocked2020 dataset^29^ and the experimental structural Binding MOAD dataset^30^.

##### CrossDocked2020^29^

For the CrossDocked2020 dataset, we followed the same filtering and splitting strategies as in previous work^12,14^, where only high-quality docking poses (RMSD < 1Å) and diverse proteins (sequence identity using MMseqs2^34^ < 30%) were retained. This resulted in 100,000 high-quality protein-ligand pairs for the training set and 100 proteins for the test set.

We ran the test set through PoseBusters^26^ and used the Open Drug Discovery Toolkit^35^ (ODDT) to check for existence of hydrogen bond interactions between the ligand and the protein. In order to ensure that the test set was of high quality (we kept only those that passed all PoseBusters tests) and that each test case contained relevant pharmacophores (we selected only examples that had more than 3 hydrogen bond interactions).

This process yielded a set of 73 complexes from the initial 100. For a full list of these complexes, please refer to section S1 in the Supplementary Information (SI).

For measuring the success rate (see the Evaluation section), we further filtered the remaining 73 complexes using the NIH filter, resulting in a final set of 48 complexes to evaluate the ‘success rate’.

##### Binding MOAD^30^

Is a dataset of experimentally determined protein-ligand complexes, we filtered and categorized based on the proteins’ enzyme commission number, following the approach by Schneuing et al.^16^ This yields 40,344 protein-ligand pairs for training and 130 pairs for testing.

Similar to the CrossDocked2020 dataset, our selection criteria for the test set involved passing all PoseBusters tests^26^ and having more than 3 hydrogen bonds. This yielded a set of 43 complexes from the initial 130. For a detailed list of these complexes, please refer to section S2 in the SI.

#### 3.1.2 Comparison to other methods

We compared MolSnapper against other methods designed to condition ligand generation within a protein pocket.

##### MolDiff

To validate the effectiveness of our conditioning, we compared it to MolDiff, without conditioning. Following molecule generation, the ligands generated by MolDiff were docked using the Vina method^36^ as molecules generated by MolDiff are not built in the pocket.

##### SILVR^25^

is a method tailored for generating molecules to fit into protein binding sites. We utilized their code, specifying pharmacophore positions and atom types as references, treating the remaining atoms as dummy atoms. The SILVR rate was set to 0.01, following the original paper.

##### DiffSBDD^16^

employs Inpainting, a conditioning method for flexible molecule design. For DiffSBDD, we provided pharmacophore positions and atom types as input for the fixed atoms.

In all experiments, we generated 100 new ligands for each reference, allowing three attempts for each generation.

#### 3.1.3 Evaluation

##### Shape and color similarity score (SC_RDKit_)

assesses the 3D similarity between generated molecules and a reference molecule, as described in^22^. SC_RDKit_ scores range from 0 (no match) to 1 (perfect match). The color similarity function evaluates the overlap of pharmacophoric features, while the shape similarity measure involves a simple volumetric comparison between the two conformers. SC_RDKit_ uses two RDKit functions, based on the methods described in Putta et al.^37^ and Landrum et al.^38^.

##### Synthetic Accessibility (SA)

measures how easily a compound can be synthesized. It assesses molecular complexity and leverages historical data from synthesized chemicals to estimate synthetic knowledge. High SA scores denote compounds that are simpler to synthesize, favored in drug development, while low SA scores highlight potential synthetic challenges^39^.

##### Same ODDT Interaction

This metric calculates the percentage of hydrogen bonds shared between the generated ligands and the protein, and the reference ligand and the protein, using the Open Drug Discovery Toolkit^35^ (ODDT).

##### PoseBusters Pass Rate

This metric indicates the percentage of generated molecules that pass PoseBusters^26^. Our evaluation focuses on ligands that pass the PoseBusters tests. In order to ensure that we only consider ligands that are physically plausible. The PoseBusters test suite examines aspects of chemical validity and consistency, intramolecular validity, and intermolecular validity, assessing physicochemical consistency and structural plausibility in the generated poses.

##### Success Rate

This metric evaluates the proportion of molecules that not only pass the initial filtering stages but also are considered more likely to be viable drug candidates. First, we exclude molecules flagged by the NIH^40,41^ filter from the test set to remove potentially problematic structures. The NIH filter, implemented using RDKit, screens out substructures known for undesirable properties such as reactive functionalities and medicinal chemistry exclusions. We then apply thresholds for SA and QED based on the minimum values found in the test set. We report the mean number of generated molecules that, after passing PoseBusters, also passed this filtering, in addition to reporting the number of pockets without ligands at all.

As part of our evaluation strategy, we employed a focused approach centered on SC_RDKit_ scores. For each metric (SA and Same ODDT Interaction), we first identify the subset of ligands with the best SC_RDKit_ scores – ‘Top 1’ being the single best, and ‘Top 3’ encompassing the three best-scoring ligands. Once these subsets are determined based on their SC_RDKit_ scores, we then calculate the other metrics specifically for these groups. For example, when we refer to ‘Top 3 SA,’ this is the best Synthetic Accessibility score obtained from the set of the top three ligands as ranked by their SC_RDKit_ scores. Additionally, following common practice, we excluded water molecules from the generation and evaluation process. By centering our analysis on the top SC_RDKit_ performers, we aimed to highlight those candidates most likely to succeed in real-world drug development scenarios.

### 3.2 Results and discussion

#### Conditioning methods without pocket-specific training

In table 1 we compare MolSnapper to MolDiff without any conditioning and to SILVR, a conditioning method that, like our approach, was trained on a molecule dataset alone rather than on protein-molecule data. On the CrossDocked data, SILVR only generated at least three molecules (out of 100 attempts) that passed the PoseBusters checks for 68 out of 73 complexes. Table 1 shows the results for these 68 complexes. The results for the full 73 complexes for MolSnapper and MolDiff can be found in Table S1 in the SI. The ‘success rate’ metric was determined using QED and SA thresholds of 0.19 and 0.33, respectively, based on the minimum values observed in the test set.

**Table 1.** Comparison on the CrossDocked dataset of MolDiff (without conditioning), SILVR, DiffSBDD (trained on CrossDocked) and our method, MolSnapper. Arrows next to metrics indicate superiority (up: larger is better, down: smaller is better). The best results are highlighted in bold. ‘Pass Rate’ refers to the percentage of ligands passing the PoseBusters test. ‘SC_RDKit_’ similarity to the ground truth ligand. ‘SA’ stands for Synthetic Accessibility. ‘Same ODDT Interaction’ assesses the similarity in hydrogen bonding interactions compared to a reference. ‘Success Rate’ is the proportion of molecules that pass both PoseBusters and the additional filtering criteria based on NIH, SA, and QED thresholds. ‘# Empty Pockets’ denotes the number of pockets for which no generated ligands passed the filtering criteria

| Method | Pass Rate↑ | SC<sub>RDKit</sub> ↑<br>Top 1 | SC<sub>RDKit</sub> ↑<br>Top 3 | SA ↑<br>Top 1 | SA ↑<br>Top 3 | Same ODDT<br>interaction ↑ Top 1 | Same ODDT<br>interaction ↑ Top 3 | Success Rate ↑ | # Empty<br>Pockets ↓ |
| --- | --- | --- | --- | --- | --- | --- | --- | --- | --- |
| MolDiff<br>(wo conditioning) | **91±15%** | 0.417 ± 0.111 | 0.417 ± 0.111 | **0.866 ± 0.100** | **0.907 ± 0.065** | 0.276 ± 0.217 | 0.426 ± 0.254 | 86.68 ± 12.36 | 0 |
| SILVR | 27±15% | 0.586 ± 0.148 | 0.586 ± 0.148 | 0.543 ± 0.089 | 0.610 ± 0.079 | 0.571±0.276 | 0.718 ± 0.270 | 7.05± 4.75 | 2 |
| MolSnapper | 58±22% | **0.721 ± 0.116** | **0.721 ± 0.116** | 0.631 ± 0.184 | 0.696 ± 0.163 | **0.746±0.249** | **0.810 ± 0.207** | 45.00 ± 24.35 | 2 |

Table 1 shows that MolDiff (the unconditional base model) produced ligands that have high Synthetic Accessibility (SA) scores (average 0.866 for Top 1). In order to check how these molecules may interact with the protein we docked them into the protein pocket (as described in the experiments section earlier). These docked ligands tend to pass the PoseBusters tests (91% Pose-Busters pass rate) and have a high success rate of 86.68%. However, in general, the ligands generated by MolDiff failed to recapitulate the interactions shown by the reference ligand (average same ODDT interaction 0.276 for Top 1). SILVR and MolSnapper (our method) generate ligands conditioned on the 3D pharmacophore positions and both methods generate molecules with lower average SA scores compared to unconditioned MolDiff with the MolSnapper outputs being more synthesizable (SILVR’s average Top 1 SA is 0.543 vs. MolSnapper’s 0.631). SILVR and MolSnapper also show lower PoseBusters Pass Rates than original MolDiff, however, 58% of MolSnappers molecules pass compared to only 27% of SILVRs. Furthermore, MolSnapper demonstrates a higher success rate, achieving 45% compared to SILVR’s 7% on average. The SC_RDKit_ scores of the generated molecules from MolSnapper (average 0.721 for Top 1) are significantly higher than those from MolDiff or SILVR (averages of 0.417 and 0.586 for the Top 1, respectively). Moreover, both SILVR and MolSnapper, due to their conditioning approach, better regenerate the original hydrogen bonds compared to MolDiff. MolSnapper outperforms SILVR, with an average of 0.746 for the Top 1 in the ‘Same ODDT interaction’ metric compared to 0.571 by SILVR. These results demonstrate that the MolSnapper method consistently outperforms SILVR across various metrics.

The differences in performance can be attributed to several issues observed with SILVR’s generated molecules. One issue is the presence of numerous disconnected fragments in SILVR’s generated molecules. In our evaluation, we addressed this by selecting the largest fragment generated. Additionally, clashes with the protein were observed due to SILVR’s model not incorporating protein information. This is further compounded by the fact that our base model, MolDiff^15^, surpasses the base model that was used for SILVR, EDM^9^, primarily because it models and diffuses the bonds of the molecule, resulting in the generation of molecules with better validity and Synthetic Accessibility^42^.

Figure 3 provides examples of generated molecules from the SILVR and MolSnapper methods, focusing on the best SC_RDKit_-scoring ligands. In the first row, SILVR struggles to generate a molecule satisfying the given pharmacophore constraint, with the relatively small fragment unable to form a hydrogen bond with the protein. In the second row, a molecule of an appropriate size is generated, but it fails to form the same bonds as the reference ligand.

**Fig. 3.**
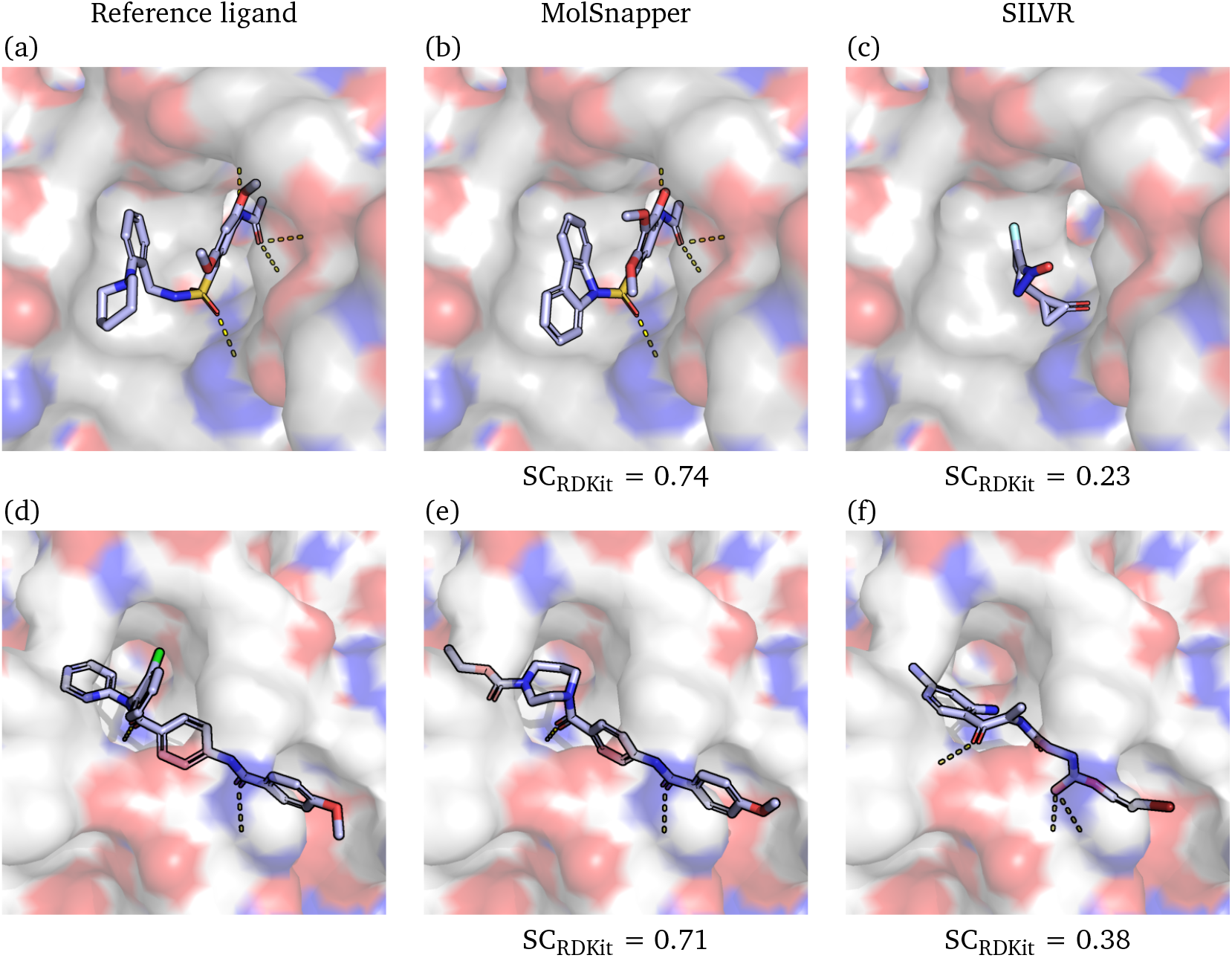
Examples from the CrossDocked dataset. For the first row: (a) Reference ligand 4aaw_A_rec_4ac3_r83_lig_tt_min_0, (b) Ligand generated by MolSnapper, (c) Ligand generated by SILVR. For the second row: (d) Reference ligand 5aeh_A_rec_5aeh_8ir_lig_tt_docked_0, (e) Ligand generated by MolSnapper, (f) Ligand generated by SILVR. The corresponding SC_RDKit_ scores are indicated below the respective images. Protein surface is colored by electrostatic potential: red for acidic, blue for basic, and white for neutral areas. Ligands are shown in stick representation. Hydrogen bonds are depicted as dashed yellow lines.

Our evaluation of MolSnapper and SILVR does not involve redocking, a common practice aimed at refining initially generated poses^27^. Redocking, while potentially useful, does not preserve the information from the pose generated by the conditioning method and therefore negates the point of a conditioned model. The authors of PoseCheck^27^ demonstrated both that the generated molecule poses are changed and that generated molecules, when redocked, can mask nonphysical features, for instance, steric clashes, hydrogen placement issues, and high strain energies.

Table S1 in the SI shows that the conservation of hydrogen bonds, as measured by ‘Same ODDT Interaction’, decreases approximately by 30% when redocking is carried out. The average RMSD of our top SC_RDKit_ generated molecules before and after docking is ∼3.44Å, with a standard deviation of ∼0.97Å.

Without redocking, MolSnapper achieves a 58% PoseBusters Pass Rate, high similarity to the original ligand, and the regeneration of most existing hydrogen bond interactions. The importance of these aspects lies in the preservation of crucial interactions and similarity to the real ligand, which are pivotal for maintaining the pharmacological relevance and effectiveness of the generated molecules.

#### Conditioning methods with pocket-specific training

We also benchmarked our method against the inpainting conditioning approach employed by DiffSBDD^16^, a 3D-conditional diffusion model specifically trained on protein-ligand complex data. In our comparison, we evaluated MolSnapper against DiffSBDD trained on CrossDocked data using the CrossDocked dataset where the ‘success rate’ metric was determined using QED and SA thresholds set at 0.19 and 0.33. Additionally, to ensure a more comprehensive assessment that better simulates real-world drug discovery challenges, we extended our analysis to a more demanding dataset aligning with actual drug discovery—the Binding MOAD dataset. Here, we compared MolSnapper to DiffSBDD trained on Binding MOAD where the QED and SA thresholds were set at 0.22 and 0.31..

Table 2 illustrates that DiffSBDD and MolSnapper show comparable performance in the generation of molecules that are similar to the original molecule (SC_RDKit_) and in recapitulating the binding interactions (same ODDT interactions). However, unlike DiffSBDD, MolSnapper does not require a dataset of complexes for training, using the same model for both test sets. Additionally, MolSnapper produces both more physically viable and drug-like molecules (PoseBuster pass rate and success rate) and more Synthetically Accessible ones (SA score). These results indicate that training on the larger molecule space and conditioning for pocket generation rather than training only on the protein-molecule set gives a wider and therefore a better representation of real molecules.

**Table 2.** Comparison on the CrossDocked dataset between DiffSBDD (trained on CrossDocked) and our method, MolSnapper, and on Binding MOAD between DiffSBDD (trained on Binding MOAD) and MolSnapper. Arrows next to metrics indicate superiority (up: larger is better, down: smaller is better). The best results are highlighted in bold. ‘Pass Rate’ refers to the percentage of ligands passing the PoseBusters test. ‘SC_RDKit_’ similarity to the ground truth ligand. ‘SA’ stands for Synthetic Accessibility. ‘Same ODDT Interaction’ assesses the similarity in hydrogen bonding interactions compared to a reference. ‘Success Rate’ is the proportion of molecules that pass both PoseBusters and the additional filtering criteria based on NIH, SA, and QED thresholds. ‘# Empty Pockets’ denotes the number of pockets for which no generated ligands passed the filtering criteria

| Data | Method | Pass Rate↑ | SC<sub>RDKit</sub> ↑<br>Top 1 | SC<sub>RDKit</sub> ↑<br>Top 3 | SA ↑<br>Top 1 | SA ↑<br>Top 3 | Same ODDT<br>interaction ↑ Top 1 | Same ODDT<br>interaction ↑ Top 3 | Success Rate ↑ | # Empty<br>Pockets ↓ |
| --- | --- | --- | --- | --- | --- | --- | --- | --- | --- | --- |
| CrossDocked | DiffSBDD (CrossDocked) | 47±19% | 0.628 ± 0.141 | 0.628 ± 0.141 | 0.638 ± 0.118 | 0.695 ± 0.094 | **0.783 ± 0.219** | **0.874 ± 0.160** | 24.6± 19.13 | 1 |
|  | MolSnapper | **58±22%** | **0.714 ± 0.116** | **0.714 ± 0.116** | **0.642 ± 0.182** | **0.722 ± 0.162** | 0.746 ± 0.246 | 0.824 ± 0.204 | **44.58 ± 24.28** | 2 |
| Binding MOAD | DiffSBDD (Binding MOAD) | 31±16% | **0.557±0.146** | **0.557±0.146** | 0.545±0.125 | 0.601 ± 0.093 | 0.839 ± 0.321 | 0.938 ± 0.182 | 13.26 ± 8.05 | 0 |
|  | MolSnapper | **57±18%** | 0.537 ± 0.075 | 0.537 ± 0.075 | **0.706 ± 0.109** | **0.784 ± 0.076** | **0.937 ± 0.221** | **0.968 ± 0.160** | **33.56 ± 21.99** | 0 |

#### Structure-based drug design for a specific target

In this section, we further demonstrate the applicability of MolSnapper through two study cases from the literature. These cases involve drug development where either a shared pharmacophore with inhibitors was targeted or there was a need to mimic interactions known from existing inhibitors. The second case highlights MolSnapper’s capability for scaffold hopping, showing that it can be effectively used not just for generating whole molecules, but also for modifying and expanding existing scaffolds to achieve desired interactions.

#### Case Study 1

##### De Novo Design with Pharmacophore Constraints

Yan et al.^43^ investigated the development of metallo-*β*-lactamase (MBL) inhibitors. *β*-Lactam antibiotics, known for disrupting the integrity of bacterial cell walls, are among the most widely used drugs for treating bacterial infections. However, their efficacy is increasingly compromised by bacterial resistance mechanisms, including the production of serine-*β* -lactamases (SBLs) and metallo-*β*-lactamases (MBLs). Yan et al. designed 2-aminothiazole-4-carboxylic acids (AtCs) (PDB: 8HX5) to mimic the binding interactions of carbapenem hydrolysate with MBLs (PDB: 6Y6J). By introducing 2-aminothiazole-4-carboxylic acid as a core scaffold, they conducted thorough investigations to enhance potency across multiple MBL subclasses, closely mimicking the essential binding features of known MBL inhibitors, although the binding positions were not fully aligned with those of the reference inhibitor.

We applied MolSnapper using the reference inhibitor interactions targeted by Yan et al. as the primary design goal, without involving scaffold exploration. We generated 5000 molecules using MolSnapper and applied a series of filters, including QED, SA score, and RDKit filters such as PAINS^44^, Brenk^45^, and NIH^40,41^ filters. After filtering, 3061 molecules remained. Next, we checked the generated molecules with PoseBusters, refining the set further to 2716 molecules, of which 1139 were unique.

We then evaluated these molecules based on their Vina docking scores to identify the best candidates. Figure S1 in the SI shows the best Vina scores generated molecules alongside their interactions, illustrating how these generated molecules compare with the reference inhibitor.

Among the 1139 unique molecules, 55.5% exhibited the same or greater interactions as the developed compound by Yan et al. (measured with ODDT) with an overall 98% showing more interactions. In addition, 18% of these molecules had Vina docking scores better than the developed compound. We also examined the structural similarity of the generated molecules to the developed compound, despite the reference points not being fully aligned with it, and found that 125 ligands had SC_RDKit_ scores greater than 0.5. This demonstrates that MolSnapper successfully generated ligands similar to the developed compound, with comparable interactions. In Figure 4, the best-scoring molecules based on SC_RDKit_ are presented, along with a comparison of interactions between the developed compound and MolSnapper-generated molecules. Lastly, generating with SILVR method showed that 1376 molecules passed the QED, SA, and RDKit filtering. After applying PoseBusters, 920 molecules remained, with 752 unique, but none had SC_RDKit_ scores greater than 0.5, and none had Vina scores better then the developed compound.

**Fig. 4.**
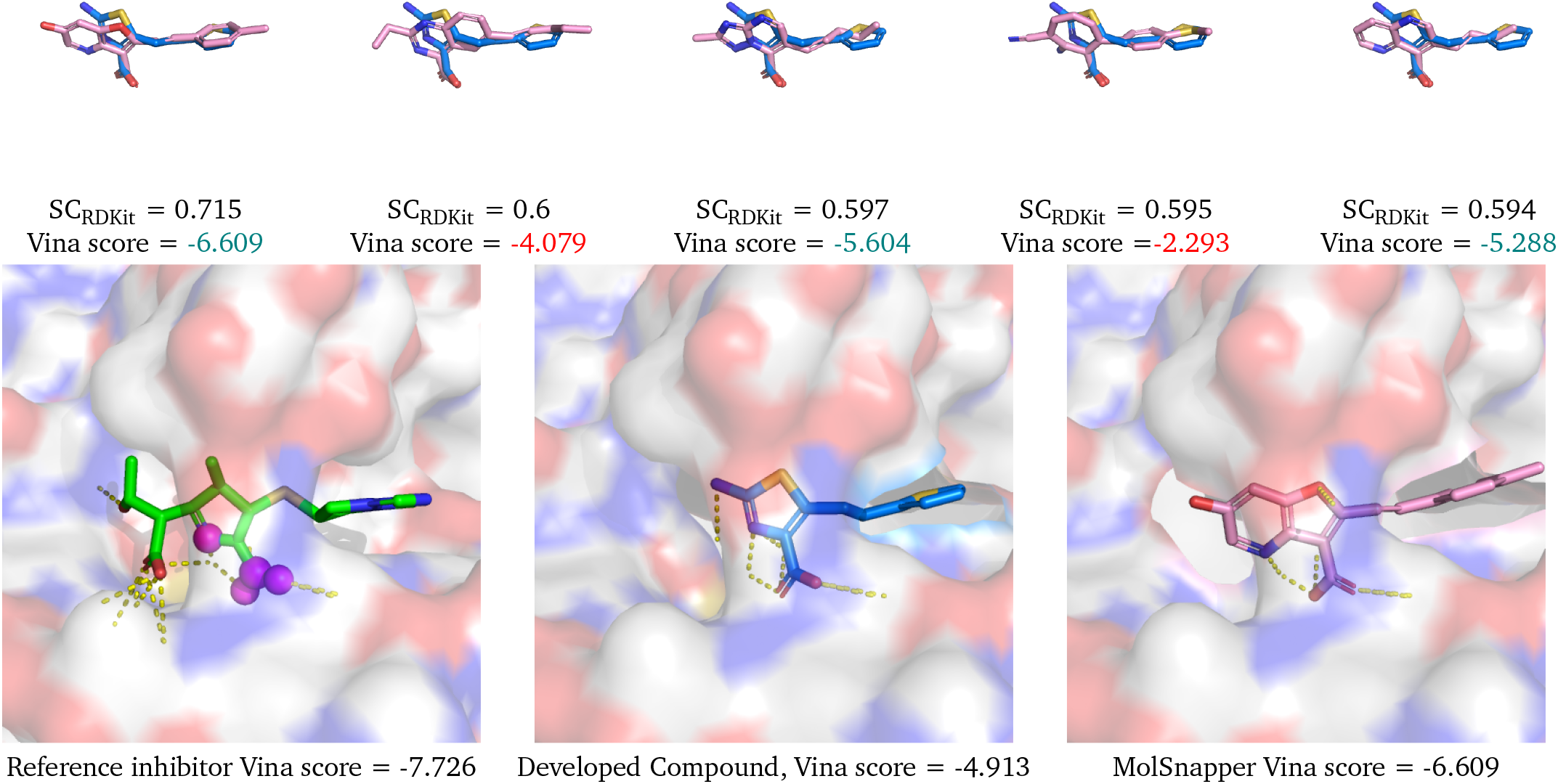
Generated molecules with the highest SC_RDKit_ scores for for Case Study 1: De Novo Design with Pharmacophore Constraints. The blue structure represents the reference compound, while the pink structures are the top molecules generated by MolSnapper. From left to right, the molecules are ordered by decreasing SC_RDKit_ score. Vina scores are also displayed; they are shown in green if better than the reference compound and in red if worse. The second row displays the reference structure in green, showing the magenta spheres indicating the interaction sites intended to be mimicked, the compound developed by Yan et al. and the most similar molecule generated by MolSnapper. Dashed yellow lines indicate the interactions between the target and the ligand.

#### Case Study 2

##### Scaffold Growing

The COVID-19 pandemic, caused by SARS-CoV-2, has led to millions of deaths and continues to challenge global health. Despite the rapid development of vaccines, there is still an urgent need for effective oral antiviral drugs to combat the virus and its variants.

Key interactions observed in the Moonshot compounds^46,47^ led to the identification of a promising drug candidate developed by Yuto Unoh et al.^48^(PDB: 7VTH). In their study, the focus was on optimizing a core scaffold to enhance binding affinity and specificity, while retaining essential interactions within the binding pocket and modifying the side chains to improve drug efficacy.

We adopted a similar approach, concentrating on the same core scaffold utilized in their research, with the goal of growing it to achieve the necessary interactions for effective inhibition. Using MolSnapper, we generated 5000 molecules and applied a series of filters, including QED, SA score, NIH, and PAINS, which reduced the set to 2413 molecules. From these, 1234 passed PoseBusters, with 1077 being unique. 49% of the unique molecules recapitulated the exact interactions observed in the reference ligand (to the same residue), while 77% demonstrated equal or additional interactions (not necessarily to the same residues). In addition, 16% of these molecules had Vina docking scores better than the reference compound.

In Figure 5, we present the top 5 generated molecules that most closely resemble the reference structure. All of these molecules share the same key interactions as the reference compound. This case study illustrates the flexibility of MolSnapper, demonstrating that it can be effectively used not only for generating whole molecules but also for scaffold growth, allowing for the refinement and enhancement of existing scaffolds to achieve desired interactions.

**Fig. 5.**
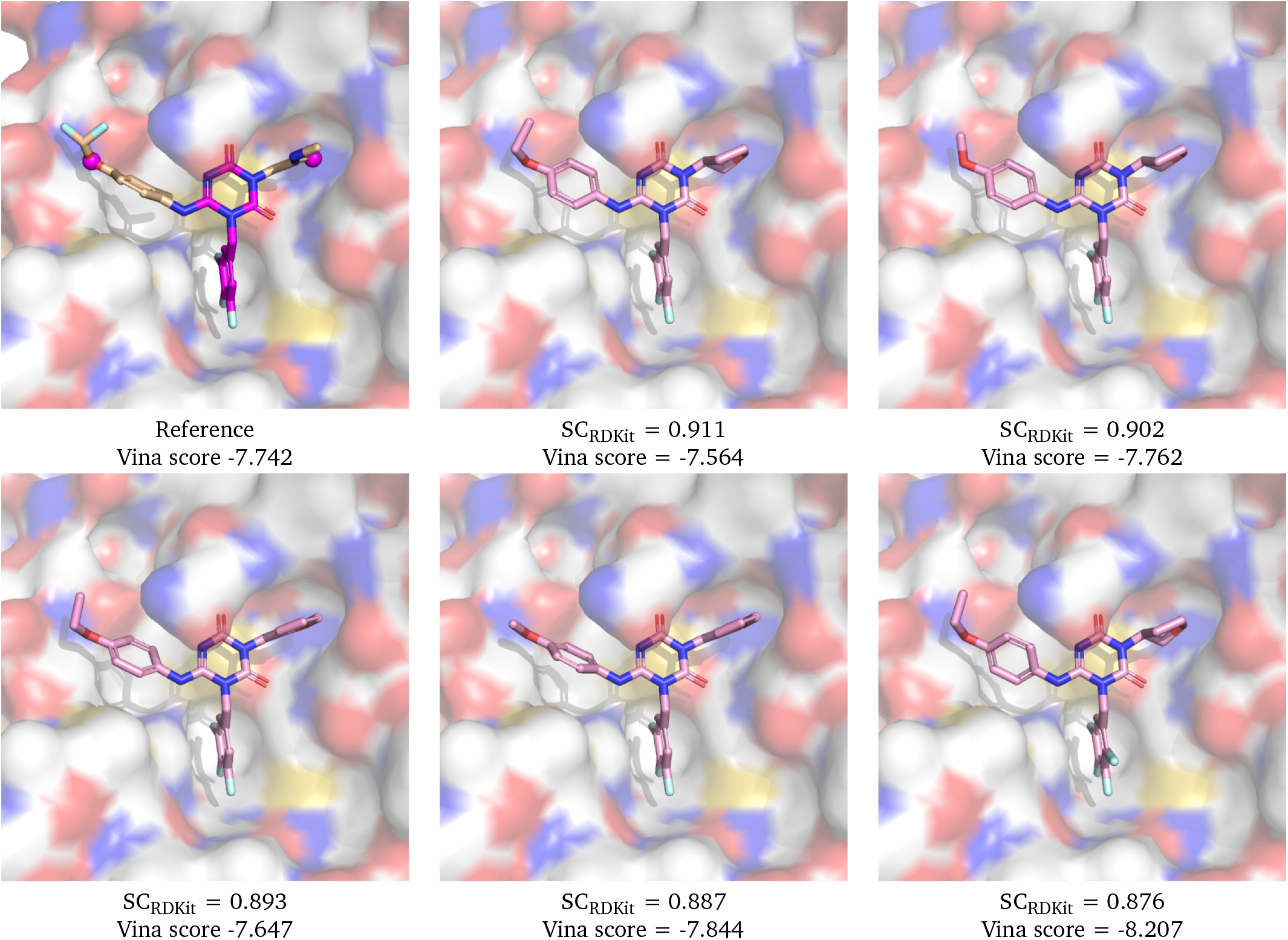
Generated molecules with the highest SC_RDKit_ scores for the COVID-19 3CLpro compound scaffold growing case study.The hit compound is shown on the left, with the main scaffold and reference points for MolSnapper generation (in magenta). The pink structures are the top SC_RDKit_ score molecules generated by MolSnapper. From left to right, the molecules are ordered by decreasing SC_RDKit_ score.

### 3.3 Ablation Studies

#### Influence of Initialization and Reference Conditioning on Generation

We compared three variations of initialization strategies and the treatment of reference points on the CrossDocked dataset.

1. Random Initialization around Pharmacophore Fixed Point with Fixed Reference Positions: The generated molecule’s initial positions are random noise around the pharmacophore fixed points and the reference positions are fixed during the diffusion reverse process.
2. Random Initialization with Fixed Reference Positions: The generated molecule’s initial positions are random noise and the reference positions remain fixed throughout the generation process as in the previous scenario.
3. Random Initialization around Pharmacophore Fixed Point with Noised Reference Points: The generated molecule’s initial positions are random noise around the pharmacophore fixed points and the reference points are noised based on a predefined noise scaling schedule during the diffusion reverse process.

Table 3 summarizes our ablation study results, focusing on the impact of noise in molecular generation using GNNs. Optimal SC_RDKit_ scores were achieved with initialization 1, random initialization around fixed pharmacophore points, but without adding noise to reference points. This suggests the importance of stable reference points in maintaining key ligand-target interactions during the generation process.

**Table 3.** Ablation studies comparing three distinct strategies for initializing and conditioning molecular generation in the CrossDocked dataset.

| Method's configuration | Pass Rate↑ | $SC_{RDKit} \uparrow$<br>Top 1 | $SC_{RDKit} \uparrow$<br>Top 3 | SA ↑<br>Top 1 | SA ↑<br>Top 3 | Same ODDT<br>interaction ↑ Top 1 | Same ODDT<br>interaction ↑ Top 3 |
| --- | --- | --- | --- | --- | --- | --- | --- |
| (1) Pharma Init. + Fixed Ref. Points | **58%** | **0.714 ± 0.116** | **0.714 ± 0.116** | **0.642 ± 0.182** | 0.722 ± 0.162 | **0.746 ± 0.246** | 0.824 ± 0.204 |
| (2) Random Init. + Fixed Ref. Points | **58%** | 0.709 ± 0.108 | 0.709 ± 0.108 | 0.638 ± 0.1180 | **0.735 ± 0.157** | 0.740 ± 0.245 | **0.842 ± 0.197** |
| (3) Pharma Init. + Noised Ref. Points | 54% | 0.705 ± 0.102 | 0.705 ± 0.102 | 0.622 ± 0.208 | 0.715 ± 0.164 | 0.736 ± 0.243 | 0.809 ± 0.220 |

## 4 Conclusions

We present MolSnapper, a novel method for conditioning diffusion models to incorporate 3D pharmacophoric constraints, offering a tool for molecular generation with controlled design processes. MolSnapper enables the easy utilization of prior knowledge to improve molecule design.

MolSnapper outperforms competing methods like SILVR and DiffSBDD, producing up to 3 times more molecules that pass Pose-Busters checks. It also offers up to a 20% improvement in the shape and color similarity to the reference ligands, leading to a 30% better retrieval of initial hits.

Our work bridges a crucial gap as it enables the use of models trained on large datasets of molecules to be conditioned for use in pocket binding. Our approach generates ligands in a controlled and effective generative process, integrating 3D structural information and expert knowledge.

## Author Contributions

YZ developed the methods and wrote the text, with input and support from CMD and BM. All authors reviewed and edited the manuscript.

## Acknowledgements

The authors thank Ruben Sanchez, Matteo Ferla, Fergus Imrie and Leo Klarner from the Oxford Protein Informatics Group for their helpful comments and discussions.

## Supplementary Information

### S1 Filtered Ground Truth Ligands from CrossDocked test dataset

1. 2z3h_A_rec_1wn6_bst_lig_tt_docked_3
2. 4aaw_A_rec_4ac3_r83_lig_tt_min_0
3. 4yhj_A_rec_4yhj_an2_lig_tt_min_0
4. 14gs_A_rec_20gs_cbd_lig_tt_min_0
5. 4rn0_B_rec_4rn1_l8g_lig_tt_min_0
6. 1fmc_B_rec_1fmc_cho_lig_tt_docked_1
7. 3daf_A_rec_3daf_feg_lig_tt_docked_0
8. 5w2g_A_rec_5w2i_adp_lig_tt_min_0
9. 3dzh_A_rec_3u4i_cvr_lig_tt_docked_0
10. 3g51_A_rec_3g51_anp_lig_tt_min_0
11. 2jjg_A_rec_2jjg_plp_lig_tt_min_0
12. 2rhy_A_rec_2rhy_mlz_lig_tt_min_0
13. 2pqw_A_rec_2rhy_mlz_lig_tt_min_0
14. 5bur_A_rec_5x8f_amp_lig_tt_docked_7
15. 3gs6_A_rec_2oxn_oan_lig_tt_docked_4
16. 1r1h_A_rec_1r1h_bir_lig_tt_docked_1
17. 1dxo_C_rec_1gg5_e09_lig_tt_min_0
18. 1gg5_A_rec_1kbo_340_lig_tt_min_0
19. 5b08_A_rec_5b09_4mx_lig_tt_min_0
20. 2azy_A_rec_2azy_chd_lig_tt_docked_0
21. 5i0b_A_rec_5vef_m77_lig_tt_min_0
22. 1phk_A_rec_1phk_atp_lig_tt_min_0
23. 4keu_A_rec_4ket_pg4_lig_tt_min_0
24. 1djy_A_rec_1djz_ip2_lig_tt_min_0
25. 5l1v_A_rec_5l1v_7pf_lig_tt_docked_0
26. 2rma_A_rec_3rdd_ea4_lig_tt_docked_0
27. 4p6p_A_rec_4p77_5rp_lig_tt_docked_0
28. 3u5y_B_rec_3u57_dh8_lig_tt_min_0
29. 4f1m_A_rec_4f1m_acp_lig_tt_min_0
30. 4tqr_A_rec_2xca_doc_lig_tt_min_0
31. 4lfu_A_rec_4y13_480_lig_tt_min_0
32. 3jyh_A_rec_3n0t_opy_lig_tt_min_0
33. 4iwq_A_rec_4jlc_su6_lig_tt_min_0
34. 1l3l_A_rec_1l3l_lae_lig_tt_min_0
35. 1e8h_A_rec_1e8h_adp_lig_tt_min_0
36. 2e24_A_rec_1j0n_ceg_lig_tt_docked_14
37. 2hcj_B_rec_2hcj_gdp_lig_tt_docked_0
38. 3kc1_A_rec_3kc1_2t6_lig_tt_min_0
39. 4ja8_B_rec_4ja8_1k9_lig_tt_docked_0
40. 4iiy_A_rec_3tle_gsu_lig_tt_docked_12
41. 3v4t_A_rec_4e7f_udp_lig_tt_min_0
42. 3tym_A_rec_3n5v_xfh_lig_tt_min_0
43. 4d7o_A_rec_3n5z_xfm_lig_tt_min_0
44. 4kcq_A_rec_4cwv_hw8_lig_tt_min_0
45. 1umd_B_rec_1umb_tdp_lig_tt_docked_1
46. 4pxz_A_rec_4pxz_6ad_lig_tt_min_0
47. 2cy0_A_rec_2d5c_skm_lig_tt_min_0
48. 3w83_B_rec_2e6d_fum_lig_tt_min_0
49. 2e6d_A_rec_2e6d_fum_lig_tt_min_0
50. 4rv4_A_rec_4rv4_prp_lig_tt_docked_2
51. 5d7n_D_rec_4jt9_1ns_lig_tt_min_0
52. 4tos_A_rec_4tos_355_lig_tt_min_0
53. 5aeh_A_rec_5aeh_8ir_lig_tt_docked_0
54. 4rlu_A_rec_4rlu_hcc_lig_tt_min_0
55. 4xli_B_rec_4xli_1n1_lig_tt_min_0
56. 3l3n_A_rec_2iux_nxa_lig_tt_docked_2
57. 5tjn_A_rec_1zj1_nlc_lig_tt_docked_4
58. 5liu_X_rec_4gq0_qap_lig_tt_min_0
59. 3o96_A_rec_3o96_iqo_lig_tt_docked_2
60. 4qlk_A_rec_4qlk_ctt_lig_tt_docked_0
61. 3hy9_B_rec_3hyg_099_lig_tt_min_0
62. 4bel_A_rec_2ewy_dbo_lig_tt_min_0
63. 3nfb_A_rec_3nfb_oae_lig_tt_docked_2
64. 4m7t_A_rec_4m7t_sam_lig_tt_min_0
65. 3u9f_C_rec_3u9f_clm_lig_tt_min_0
66. 2f2c_B_rec_1xo2_fse_lig_tt_min_0
67. 3chc_B_rec_3ch9_xrg_lig_tt_min_0
68. 4z2g_A_rec_4z2g_m6v_lig_tt_docked_16
69. 3af2_A_rec_3af4_gcp_lig_tt_min_0
70. 1jn2_P_rec_1val_png_lig_tt_docked_11
71. 3li4_A_rec_2gvv_di9_lig_tt_min_0
72. 4azf_A_rec_5lxc_7aa_lig_tt_min_0
73. 2pc8_A_rec_1eqc_cts_lig_tt_docked_6
74. 7AKI-bio1_RJQ_A_202

### S2 Filtered Ground Truth Ligands from Binding MOAD test dataset

1. 2FKY-bio2_N2T_B_605
2. 3EKU-bio1_CY9_A_903
3. 1RDW-bio1_LAR_X_391
4. 5NCP-bio1_EG5_A_201
5. 7AKI-bio1_RJQ_A_203
6. 5NEG-bio1_8VK_B_201
7. 5NCG-bio1_8TB_A_202
8. 6RCJ-bio1_K0H_A_201
9. 2FL6-bio2_N5T_B_605
10. 3F7H-bio2_419_B_1
11. 3F7I-bio2_G13_B_1
12. 2I3I-bio1_618_A_501
13. 3GTA-bio2_851_B_1
14. 2Q0U-bio1_LAB_A_401
15. 2FKY-bio1_N2T_A_604
16. 2I3I-bio2_618_B_501
17. 3EKS-bio1_CY9_A_903
18. 5NCG-bio2_8TB_B_202
19. 3CJO-bio1_K30_A_1
20. 5XDU-bio1_ZI6_A_403
21. 1ESV-bio1_LAR_A_401
22. 4PA0-bio2_2OW_B_1101
23. 3GT9-bio2_516_B_1
24. 3GTA-bio1_851_A_1
25. 5NCG-bio2_8TB_B_201
26. 5NDU-bio2_8V2_B_201
27. 2A5X-bio1_LAR_A_379
28. 2Q2Y-bio1_MKR_A_604
29. 5NCG-bio1_8TB_A_201
30. 3GT9-bio1_516_A_1
31. 1SQK-bio1_LAR_A_378
32. 6RCF-bio1_K0K_A_201
33. 3F7H-bio1_419_A_1
34. 5NCF-bio2_8T5_B_204
35. 4PA0-bio1_2OW_A_1101
36. 5ZZB-bio2_LAB_D_401
37. 2FL2-bio2_N4T_B_605
38. 5NDU-bio1_8V2_A_201
39. 2Q2Y-bio2_MKR_B_605
40. 5NEG-bio1_8VK_A_201
41. 3F7I-bio1_G13_A_1
42. 3MN5-bio1_LAB_A_376

### S3 Implementaion details

In our experiments on Crossdockes and Binding MOAD we enforce that nitrogen (N) is selected for donor atoms, and oxygen (O) is chosen for acceptor atoms.

Case study 1: The PAINS filter was applied to exclude molecules likely to cause nonspecific binding and false positives, while the Brenk filter was used to remove substructures associated with poor pharmacokinetics or toxicity. Additionally, the NIH filter provided an extra layer of safety screening. The thresholds used were QED ≥ 0.25 and SA score≥ 0.5.

Case Study 2 : In this case, only the PAINS and NIH filters were applied, as the reference molecule did not pass the Brenk filter. The same thresholds were used: QED ≥ 0.25 and SA score≥ 0.5.

**Table S1:** Comparison on the CrossDocked dataset of MolDiff (without conditioning, which was docked with the Vina docking method), DiffSBDD (trained on CrossDocked), and our method, MolSnapper. Arrows next to metrics indicate superiority (up: larger is better, down: smaller is better). The best results are highlighted in bold. ‘Pass Rate’ refers to the percentage of ligands passing the PoseBusters test. ‘SC_RDKit_’ similarity to the ground truth ligand. ‘SA’ stands for Synthetic Accessibility. ‘Same ODDT Interaction’ assesses the similarity in hydrogen bonding interactions compared to a reference

| Method | Pass Rate↑ | SC<sub>RDKit</sub> ↑<br>Top 1 | SC<sub>RDKit</sub> ↑<br>Top 5 | SA ↑<br>Top 1 | SA ↑<br>Top 5 | Same ODDT<br>interaction ↑ Top 1 | Same ODDT<br>interaction ↑ Top 5 |
| --- | --- | --- | --- | --- | --- | --- | --- |
| MolDiff<br>(wo conditioning) | 92±15% | 0.421 ± 0.109 | 0.421 ± 0.109 | **0.86 ± 0.096** | **0.916 ± 0.060** | 0.284 ± 0.217 | 0.496 ± 0.262 |
| DiffSBDD (CrossDocked) | 47±19% | 0.628 ± 0.141 | 0.628 ± 0.141 | 0.638 ± 0.118 | 0.695 ± 0.094 | **0.783 ± 0.219** | **0.874 ± 0.160** |
| MolSnapper | 58±22% | **0.714 ± 0.116** | **0.714 ± 0.116** | 0.642 ± 0.182 | 0.722 ± 0.162 | 0.746 ± 0.246 | 0.824 ± 0.204 |
| MolSnapper redocked | 61±21% | **0.714 ± 0.116** | **0.714 ± 0.116** | 0.642 ± 0.182 | 0.722 ± 0.162 | 0.527 ± 0.311 | 0.716 ± 0.236 |

**Figure S1.**
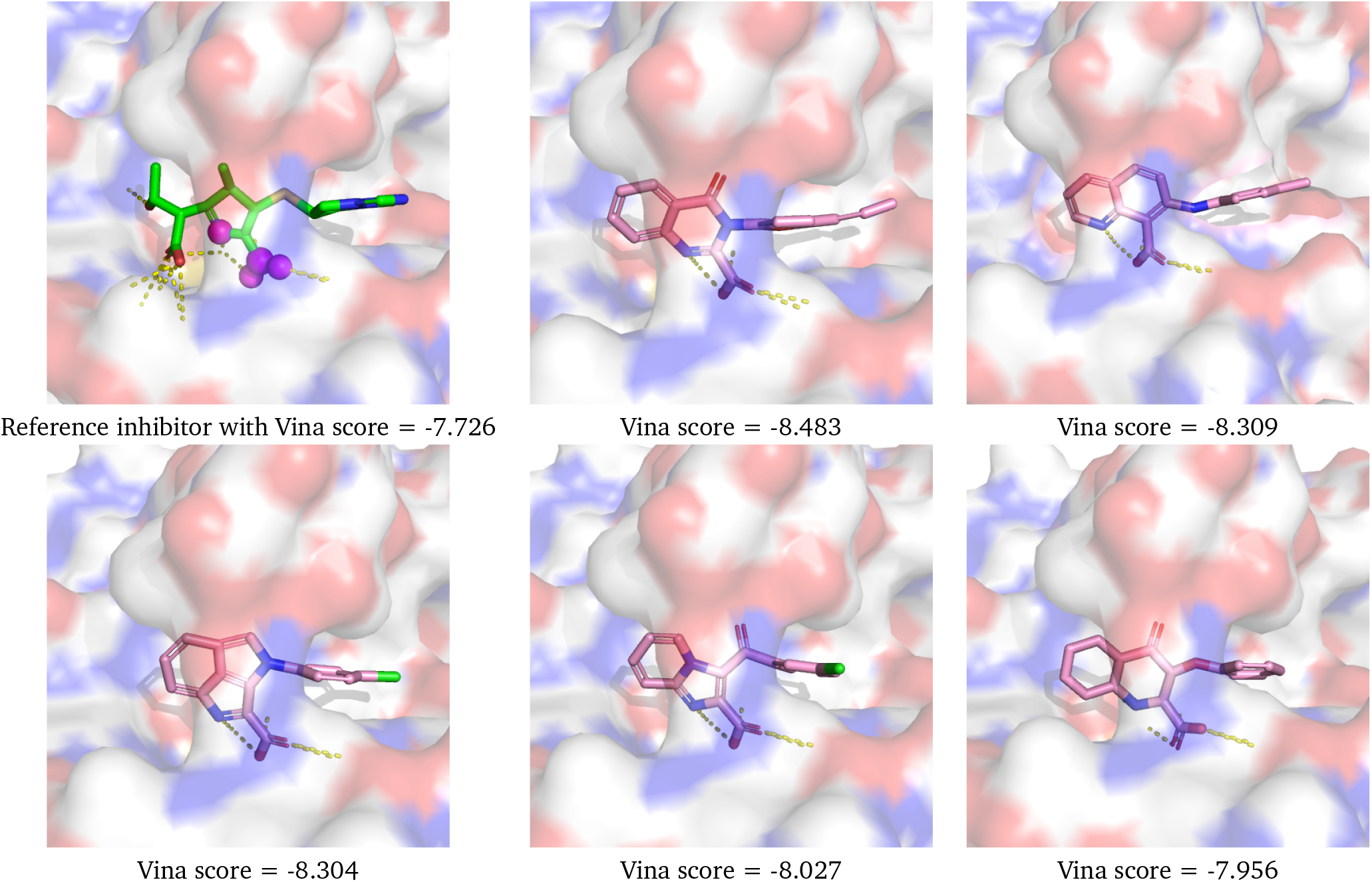
Comparison of the reference inhibitor with the top five MolSnapper-generated molecules based on Vina docking scores for Case Study 1: De Novo Design with Pharmacophore Constraints. The reference inhibitor is shown in green, with the interaction sites intended to be mimicked highlighted in magenta spheres, alongside the top five MolSnapper-generated molecules ranked by the highest Vina docking scores.

## Notes and references

1 S. M. Paul, D. S. Mytelka, C. T. Dunwiddie, C. C. Persinger, B. H. Munos, S. R. Lindborg and A. L. Schacht, Nature reviews Drug discovery, 2010, 9, 203–214.

2 J. Avorn, New England Journal of Medicine, 2015, 372, 1877–1879.

3 S.-s. Ou-Yang, J.-y. Lu, X.-q. Kong, Z.-j. Liang, C. Luo and H. Jiang, Acta Pharmacologica Sinica, 2012, 33, 1131–1140.

4 J. Ma, R. P. Sheridan, A. Liaw, G. E. Dahl and V. Svetnik, Journal of chemical information and modeling, 2015, 55, 263–274.

5 A. Mayr, G. Klambauer, T. Unterthiner and S. Hochreiter, Frontiers in Environmental Science, 2016, 3, 80.

6 A. Halevy, P. Norvig and F. Pereira, IEEE intelligent systems, 2009, 24, 8–12.

7 C. Sun, A. Shrivastava, S. Singh and A. Gupta, bProceedings of the IEEE international conference on computer vision, 2017, pp. 843–852.

8 N. Gebauer, M. Gastegger and K. Schütt, Advances in neural information processing systems, 2019, 32, year.

9 E. Hoogeboom, V. G. Satorras, C. Vignac and M. Welling, International conference on machine learning, 2022, pp. 8867–8887.

10 M. Xu, L. Yu, Y. Song, C. Shi, S. Ermon and J. Tang, International Conference on Learning Representations, 2021.

11 M. Thomas, A. Bender and C. de Graaf, Current Opinion in Structural Biology, 2023, 79, 102559.

12 S. Luo, J. Guan, J. Ma and J. Peng, Advances in Neural Information Processing Systems, 2021, 34, 6229–6239.

13 M. Liu, Y. Luo, K. Uchino, K. Maruhashi and S. Ji, International Conference on Machine Learning (ICML), 2022.

14 X. Peng, S. Luo, J. Guan, Q. Xie, J. Peng and J. Ma, International Conference on Machine Learning, 2022, pp. 17644–17655.

15 X. Peng, J. Guan, Q. Liu and J. Ma, Proceedings of the 40th International Conference on Machine Learning, 2023, pp. 27611–27629.

16 A. Schneuing, Y. Du, C. Harris, A. Jamasb, I. Igashov, W. Du, T. Blundell, P. Lió, C. Gomes, M. Welling et al., arXiv preprint arXiv:2210.13695, 2022.

17 H. Lin, Y. Huang, M. Liu, X. Li, S. Ji and S. Z. Li, arXiv preprint arXiv:2211.11214, 2022.

18 J. Guan, W. W. Qian, X. Peng, Y. Su, J. Peng and J. Ma, The Eleventh International Conference on Learning Representations, 2022.

19 J. Ho, A. Jain and P. Abbeel, Advances in neural information processing systems, 2020, 33, 6840–6851.

20 J. Guan, X. Zhou, Y. Yang, Y. Bao, J. Peng, J. Ma, Q. Liu, L. Wang and Q. Gu, 2023.

21 G. Durant, F. Boyles, K. Birchall, B. Marsden and C. Deane, bioRxiv, 2023, 2023–10.

22 F. Imrie, T. E. Hadfield, A. R. Bradley and C. M. Deane, Chemical science, 2021, 12, 14577–14589.

23 H. Zhu, R. Zhou, D. Cao, J. Tang and M. Li, Nature Communications, 2023, 14, 6234.

24 T. E. Hadfield, F. Imrie, A. Merritt, K. Birchall and C. M. Deane, Journal of Chemical Information and Modeling, 2022, 62, 2280–2292.

25 N. T. Runcie and A. S. Mey, Journal of chemical information and modeling, 2023, 63, 5996–6005.

26 M. Buttenschoen, G. M. Morris and C. M. Deane, Chemical Science, 2024.

27 C. Harris, K. Didi, A. Jamasb, C. Joshi, S. Mathis, P. Lio and T. Blundell, NeurIPS 2023 Generative AI and Biology (GenBio) Workshop, 2023.

28 D. Schaller, D. Šribar, T. Noonan, L. Deng, T. N. Nguyen, S. Pach, D. Machalz, M. Bermudez and G. Wolber, Wiley Interdisciplinary Reviews: Computational Molecular Science, 2020, 10, e1468.

29 P. G. Francoeur, T. Masuda, J. Sunseri, A. Jia, R. B. Iovanisci, Snyder and D. R. Koes, Journal of chemical information and modeling, 2020, 60, 4200–4215.

30 L. Hu, M. L. Benson, R. D. Smith, M. G. Lerner and H. A. Carlson, Proteins: Structure, Function, and Bioinformatics, 2005, 60, 333–340.

31 S. Axelrod and R. Gomez-Bombarelli, Scientific Data, 2022, 9, 185.

32 D. Chen, N. Oezguen, P. Urvil, C. Ferguson, S. M. Dann and T. C. Savidge, Science advances, 2016, 2, e1501240.

33 F. Sverrisson, J. Feydy, B. E. Correia and M. M. Bronstein, Proceedings of the IEEE/CVF Conference on Computer Vision and Pattern Recognition, 2021, pp. 15272–15281.

34 M. Steinegger and J. Söding, Nature biotechnology, 2017, 35, 1026–1028.

35 M. Wójcikowski, P. Zielenkiewicz and P. Siedlecki, Journal of cheminformatics, 2015, 7, 1–6.

36 J. Eberhardt, D. Santos-Martins, A. F. Tillack and S. Forli, Journal of chemical information and modeling, 2021, 61, 3891–3898.

37 S. Putta, G. A. Landrum and J. E. Penzotti, Journal of medicinal chemistry, 2005, 48, 3313–3318.

38 G. A. Landrum, J. E. Penzotti and S. Putta, Journal of computer-aided molecular design, 2006, 20, 751–762.

39 P. Ertl and A. Schuffenhauer, Journal of cheminformatics, 2009, 1, 1–11.

40 A. Jadhav, R. S. Ferreira, C. Klumpp, B. T. Mott, C. P. Austin, J. Inglese, C. J. Thomas, D. J. Maloney, B. K. Shoichet and A. Simeonov, Journal of medicinal chemistry, 2010, 53, 37–51.

41 R. G. Doveston, P. Tosatti, M. Dow, D. J. Foley, H. Y. Li, A. J. Campbell, D. House, I. Churcher, S. P. Marsden and A. Nelson, Organic & biomolecular chemistry, 2015, 13, 859–865.

42 X. Peng, J. Guan, Q. Liu and J. Ma, arXiv preprint arXiv:2305.07508, 2023.

43 Y.-H. Yan, T.-T. Zhang, R. Li, S.-Y. Wang, L.-L. Wei, X.-Y. Wang, K.-R. Zhu, S.-R. Li, G.-Q. Liang, Z.-B. Yang et al., Journal of Medicinal Chemistry, 2023, 66, 13746–13767.

44 J. B. Baell and G. A. Holloway, Journal of medicinal chemistry, 2010, 53, 2719–2740.

45 R. Brenk, A. Schipani, D. James, A. Krasowski, I. H. Gilbert, J. Frearson and P. G. Wyatt, ChemMedChem: Chemistry Enabling Drug Discovery, 2008, 3, 435–444.

46 A. Douangamath, D. Fearon, P. Gehrtz, T. Krojer, P. Lukacik, C. D. Owen, E. Resnick, C. Strain-Damerell, A. Aimon, P. Ábrányi-Balogh et al., Nature communications, 2020, 11, 5047.

47 C. M. Consortium, H. Achdout, A. Aimon, D. S. Alonzi, R. Arbon, E. Bar-David, H. Barr, A. Ben-Shmuel, J. Bennett, V. A. Bilenko et al., BioRxiv, 2020, 2020–10.

48 Y. Unoh, S. Uehara, K. Nakahara, H. Nobori, Y. Yamatsu, S. Yamamoto, Y. Maruyama, Y. Taoda, K. Kasamatsu, T. Suto et al., Journal of medicinal chemistry, 2022, 65, 6499–6512.

